# Vaccine-induced, high magnitude HIV Env-specific antibodies with Fc-mediated effector functions are insufficient to protect infant rhesus macaques against oral SHIV infection

**DOI:** 10.1101/2021.10.11.464024

**Authors:** Alan D. Curtis, Pooja T. Saha, Maria Dennis, Stella J. Berendam, S. Munir Alam, Guido Ferrari, Pamela A. Kozlowski, Genevieve Fouda, Michael Hudgens, Koen KA Van Rompay, Justin Pollara, Sallie R. Permar, Kristina De Paris

**Author notes:** <u>Corresponding Author</u> Kristina De Paris, PhD, University of North Carolina, Department of Microbiology, Center for AIDS Research, and Children’s Research Institute, 116 Manning Drive, Mary Ellen Jones Bldg., Rm 5004B, Chapel Hill, NC 27599-7292.

## Abstract

Improved access to antiretroviral therapy and antenatal care have significantly reduced in-utero and peri-partum mother-to-child HIV transmission. However, as breastmilk transmission of HIV still occurs at an unacceptable rate there remains a need to develop an effective vaccine for the pediatric population.

Previously, we compared different HIV vaccine strategies, intervals, and adjuvants in infant rhesus macaques to optimize the induction of HIV envelope (Env)-specific antibodies with Fc-mediated effector function. Here, we tested the efficacy of an optimized vaccine regimen against oral SHIV acquisition in infant macaques. One group of 12 animals was immunized with 1086.c gp120 protein adjuvanted with 3M-052 in stable emulsion and Modified Vaccinia Ankara (MVA) virus vector expressing 1086.c HIV Env, while the control group (n=12) was immunized only with empty MVA. The first vaccine dose was given within 10 days of birth and booster doses were administered at weeks 6 and 12.

The vaccine regimen induced Env-specific plasma IgG antibodies capable of antibody-dependent cellular cytotoxicity (ADCC) and phagocytosis (ADCP). Beginning at week 15, infants were exposed orally to escalating doses of heterologous SHIV-1157(QNE)Y173H once a week until infected. Despite the induction of strong Fc-mediated antibody responses, the vaccine regimen did not reduce the risk of infection, time to acquisition, or peak viremia compared to controls. Our results suggest that the non-neutralizing Env-specific antibodies with Fc effector function elicited by this vaccine regimen were insufficient for protection against heterologous oral SHIV infection shortly after the final immunization.

**IMPORTANCE:** Women of childbearing age are three times more likely to contract HIV infection than their male counterparts. Poor HIV testing rates coupled with low adherence to antiretroviral therapy (ART) result in a high risk of mother-to-infant HIV transmission, especially during the breastfeeding period. A preventative vaccine could curb pediatric HIV infections, reduce potential health sequalae, and prevent the need for lifelong ART in this population. The results of the current study imply that the HIV Env-specific IgG antibodies elicited by this candidate vaccine regimen, despite high magnitude of Fc-mediated effector function, but lack of neutralizing antibodies and polyfunctional T cell responses, were insufficient to protect infant rhesus macaques against oral virus acquisition.

## INTRODUCTION

The successful implementation of antiretroviral therapy (ART) for women living with HIV (WLWH) has resulted in a drastic reduction of in utero and peripartum mother-to-child transmission of HIV-1 in the last two decades. Yet, globally, between 400 to 500 infants continue to acquire HIV every day (1). The majority of these infections occur during the breastfeeding period. Limited access to ART in rural communities, HIV diagnosis late in pregnancy, gaps in linking antenatal care with postnatal mother and infant care, acute maternal infection during the breastfeeding period, and lack of ART adherence impede the prevention of HIV transmission by breastmilk (2-9). Transmission of HIV can occur throughout the entire breastfeeding period, with a cumulative risk increase with every month of breastfeeding (10-13). However, in many resource-limited countries, breastmilk remains a necessary choice for nutrition and to provide passive immunity to protect the infant against other endemic pathogens (6, 7, 14). Indeed, early weaning is associated with increased infant mortality (15-17), and the WHO recommends exclusive breastfeeding for 6-12 months for infants born to HIV-infected mothers (18). Infants born to mothers with known HIV-positive status are tested at birth and immediately started on ART if found to be infected, whereas infants who acquire HIV by breastfeeding often go undiagnosed until they develop clinical symptoms. Prolonged HIV replication prior to diagnosis may severely interfere with multiple aspects of normal immune and central nervous system development and impede immune reconstitution after ART initiation. Therefore, prevention strategies tailored to infants are needed to further reduce the risk of pediatric HIV infections.

In non-human primate (NHP) models of HIV, infection of neonatal and infant rhesus macaques with SHIV can be prevented by passive administration of broadly neutralizing HIV envelope (Env)-specific antibodies (bnAbs) (19-21). The use of bnAbs as potential prevention strategy in HIV exposed infants is supported by results from ongoing clinical trials that indicate that bnAbs (e.g., VRC01) are safe and well tolerated in human neonates (22). Clinical studies in human adults, however, demonstrated only a minimal risk reduction of HIV infection by preventative treatment with bnAbs (23, 24). Therefore, the development of an effective HIV vaccine remains a high priority for this risk group. While the induction of HIV bnAbs by vaccination remains challenging, antibodies with Fc-mediated effector function can be induced more consistently and have been associated with partial protection in multiple NHP vaccine/challenge studies (25-29) and in the human RV144 HIV vaccine trial (30). Furthermore, the protective effect of bnAbs is not due solely to their neutralization function, but also depends, at least in part, on the Fc-mediated effector functions of these bnAbs (31, 32).

Utilizing the pediatric rhesus macaque model, we previously compared different HIV vaccine modalities, immunization intervals and adjuvants to optimize the induction of HIV Env-specific IgG antibodies with Fc-mediated effector functions (33-35). Building on these results, in the current study, we tested the efficacy of an intramuscular (IM) vaccine consisting of a Modified Vaccinia Ankara (MVA) virus vector expressing transmitted/founder virus 1086.c gp120-combined with 1086.c HIV gp120 protein and 3M-052 adjuvant in stable emulsion against oral SHIV acquisition in infant macaques. Consistent with our prior findings, the vaccine induced high magnitude Env-specific antibodies in plasma with potent antibody-dependent cellular cytotoxicity (ADCC) and antibody-dependent cellular phagocytic (ADCP) function. Nonetheless, these responses did not protect infant rhesus macaques against subsequent heterologous oral SHIV challenge.

## METHODS

### Animals and sample collection

Twenty-four infant rhesus macaques (RM) were nursery-reared and housed in pairs at the California National Primate Research Center (Davis, CA). All animal procedures were approved by the UC Davis Institutional Animal Care and Use Committee. The study strictly adhered to the guidelines outlined in The *Guide for the Care and Use of Laboratory Animals* by the Committee on Care and Use of Laboratory Animals of the Institute of Laboratory Resources, National Resource Council. Peripheral blood was collected by venipuncture into EDTA-treated vacutainers and processed as described (36). Peripheral lymph node biopsies were collected at week 14 prior to initiation of oral challenges at week 15 as described (33). All experimental manipulations were performed under ketamine anesthesia (10mg/kg body weight) administered by the intramuscular (IM) route.

### Vaccines

The infants in the present study were randomly divided into 2 groups of 12 (Table 1; Figure 1). At week 0, infant RM assigned to the vaccine arm were primed IM with 2×10^8^ plaque forming units (PFU) of MVA-HIV 1086.c Env construct (in a volume of 0.25 ml divided over left and right biceps) (35) and 15 μg 1086Δ7 gp120 K160N protein mixed with 3M-052 adjuvant in stable emulsion (3M-052-SE) (34, 35) at a total dose volume of 0.5 ml, divided over the left and right quadriceps. The HIV Env 1086.c gp120-expressing MVA construct was produced as detailed elsewhere (37). In addition, infant vaccinees received 5×10^10^ viral particles (VP) of chimpanzee adenovirus (ChAdOx1.tSIVconsv239)-SIV Gag/Pol (0.25 ml IM divided over left and right gluteus) at week 0. Infants in the vaccine cohort received two successive IM booster immunizations with 1086.c gp120 protein in 3M-052-SE and MVA-HIV Env (both were the same dose as the priming immunization) and 2×10^8^ PFU MVA.tSIVconsv239 (gag/pol-expressing vector) in 0.25 ml, divided over left and right biceps) at weeks 6 and 12 (35). The ChAdOx1.tSIVconsv239 and MVA.tSIVconsv239 were kindly provided by Dr. Tomáš Hanke (Oxford University, Oxford, UK). Control infants received an empty MVA vector at weeks 0, 6, and 12 (Figure 1).

**Table 1:** Summary of study animals.

| Group | Animal ID | Sex | Age (days) at 1<sup>st</sup>.<br>Immunization | Challenges to<br>Infection | Infecting<br>Dose | Peak Viremia<br>(copies/ml) |
| --- | --- | --- | --- | --- | --- | --- |
| Mock | RM1 | Female | 8 | 17 | 1:100 | 5.1x10<sup>7</sup> |
| Mock | RM2 | Male | 8 | 3 | 1:1000 | 1.3x10<sup>8</sup> |
| Mock | RM3 | Male | 6 | 13 | 1:1000 | 2.5x10<sup>6</sup> |
| Mock | RM4 | Female | 6 | 14 | 1:100 | 7.1x10<sup>5</sup> |
| Mock | RM5 | Male | 5 | 7 | 1:1000 | 7.6x10<sup>6</sup> |
| Mock | RM6 | Female | 4 | 4 | 1:1000 | 1.3x10<sup>8</sup> |
| Mock | RM7 | Female | 10 | 8 | 1:1000 | 8.9x10<sup>6</sup> |
| Mock | RM8 | Male | 10 | 15 | 1:100 | 4.3x10<sup>7</sup> |
| Mock | RM9 | Female | 7 | 2 | 1:1000 | 2.1x10<sup>7</sup> |
| Mock | RM10 | Male | 7 | 30 | undiluted | 3.1x10<sup>4</sup> |
| Mock | RM11 | Male | 6 | 2 | 1:1000 | 1.6x10<sup>7</sup> |
| Mock | RM12 | Male | 4 | 3 | 1:1000 | 3.5x10<sup>8</sup> |
| Vaccine | RM13 | Female | 8 | 3 | 1:1000 | 5.3x10<sup>6</sup> |
| Vaccine | RM14 | Male | 7 | 15 | 1:100 | 1.7x10<sup>7</sup> |
| Vaccine | RM15 | Male | 5 | 24 | 1:10 | 5.1x10<sup>5</sup> |
| Vaccine | RM16 | Male | 5 | 1 | 1:1000 | 2.0x10<sup>6</sup> |
| Vaccine | RM17 | Male | 4 | 4 | 1:1000 | 1.1x10<sup>6</sup> |
| Vaccine | RM18 | Male | 3 | 2 | 1:1000 | 1.2x10<sup>7</sup> |
| Vaccine | RM19 | Female | 8 | 28 | 1:2 | 9.7x10<sup>6</sup> |
| Vaccine | RM20 | Male | 8 | 3 | 1:1000 | 4.7x10<sup>6</sup> |
| Vaccine | RM21 | Female | 7 | 7 | 1:1000 | 2.1x10<sup>6</sup> |
| Vaccine | RM22 | Male | 7 | 15 | 1:100 | 4.5x10<sup>7</sup> |
| Vaccine | RM23 | Female | 6 | 20 | 1:100 | 5.3x10<sup>7</sup> |
| Vaccine | RM24 | Male | 6 | 23 | 1:10 | 1.4x10<sup>7</sup> |

**Figure 1:**
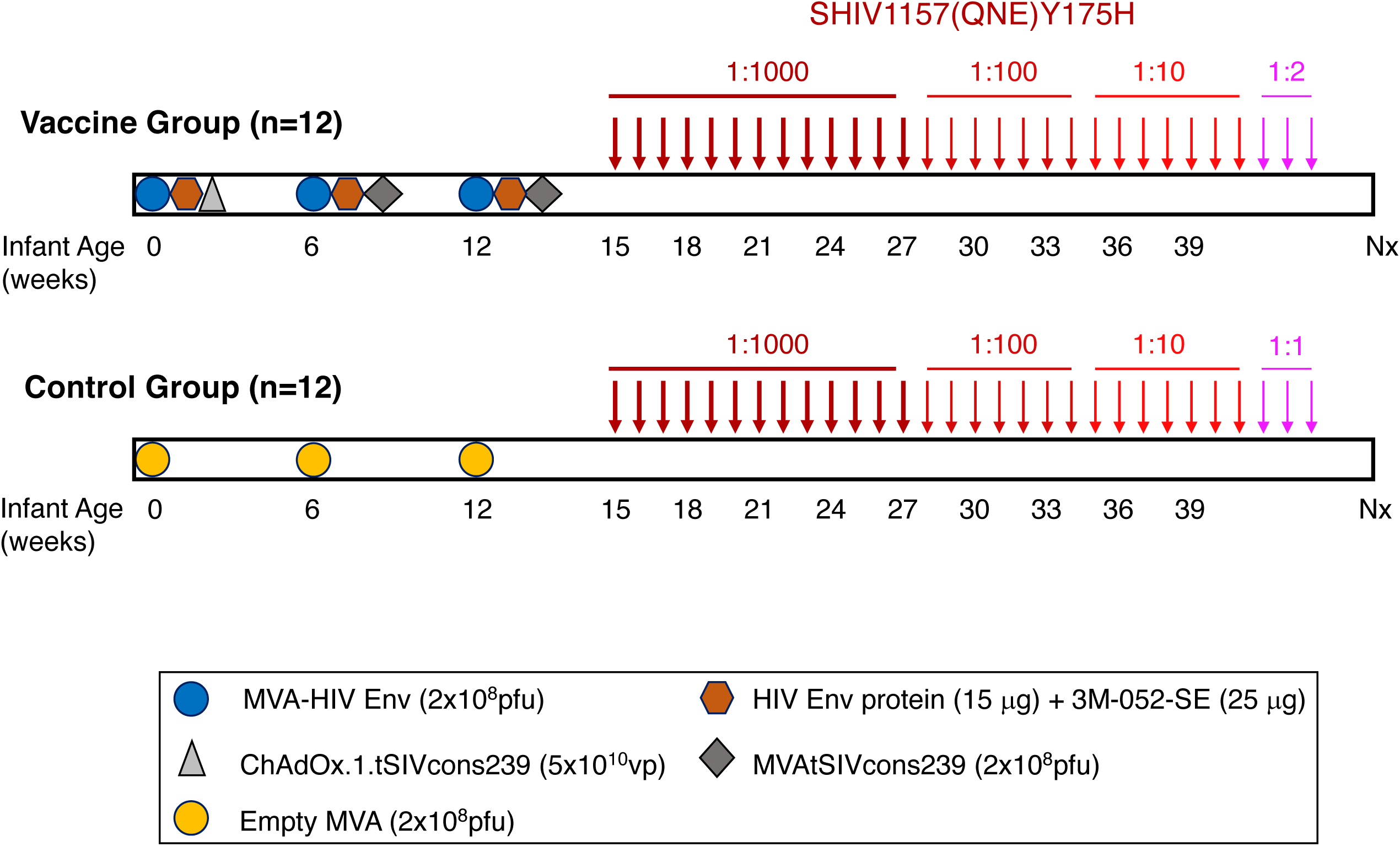
Experimental design. In the vaccine group, 12 neonatal rhesus macaques (Table 1) were immunized with 2×10^8^ PFU MVA-HIV Env (blue circle), HIV Env protein (15 μg) mixed with 3M-052-SE (brown hexagon), and 5×10^10^ ChAdOx1.tSIVconsv239 viral particles (light grey triangle) at week 0. Booster immunizations of 2×10^8^ PFU each of MVA-HIV Env, HIV Env protein in 3M-052-SE, and MVA.tSIVconsv239 (dark grey diamond) were provided at weeks 6 and 12. A second cohort of 12 age-matched RM received control MVA (orange circle) immunizations at weeks 0, 6, and 12. Beginning at week 15, animals were challenged weekly with SHIV-1157(QNE)Y173H viral stock diluted 1:1000 in RPMI until infected. After 13 exposures, uninfected infants (n=11) were exposed to a 1:100 SHIV dose for 7 weeks, a dose that was increased to 1:10 for seven more exposures in animals not infected by the 1:100 dose (n=4). Two infants remained negative and became infected after challenge with 1:2 dilution of virus stock (RM19) or undiluted (1:1) virus (RM10), respectively (Table 1). SHIV exposures are indicated by arrows with distinct shades of red based on virus dilution.

### SHIV-1157(QNE)Y173H challenge of vaccinated and control macaques

At week 15, 3 weeks after the last immunization, infant macaques were orally exposed once weekly to Tier 2 SHIV-1157(QNE)Y173H, a derivative of the CCR5-tropic clade C SHIV-1157ipd3N4 (28), that was kindly provided by Dr. Sampa Santra (Harvard University, Boston, MA). The virus stock corresponded to 3.7×10^9^ copies/ ml and had a TCID_50_ of 4.88×10^8^/ml in TZM-bl cells (28, 38). SHIV-1157(QNE)Y173H (henceforth referred to as SHIV) was selected for its high sequence homology to the 1086.c V1V2 region (28). Virus was administered as a 1:1000 dilution of virus stock in 1 ml sucrose-containing RPMI 1640 medium in a needleless syringe (39). Infants were considered to be systemically infected following two consecutive PCR-positive values (see below). After 13 challenges of 1:1000, uninfected infants (n=11) received an increased dose of 1:100. Following 7 challenges with 1:100 diluted virus, the viral challenge was increased to 1:10 dilution in the remaining uninfected animals (n=4). Two infants (RM19 and RM10) remained negative and became infected after challenge with 1:2 dilution of virus stock or undiluted virus, respectively (Table 1). Approximately 12 weeks post SHIV infection, animals were euthanized.

### SHIV RNA quantification

Weekly quantitative analysis of SHIV RNA in plasma began on week 16 as previously described (36). Briefly, RNA was manually extracted from limited plasma volumes and assayed by reverse transcription-PCR (RT-PCR) with a limit of detection of 15 copies/ ml. Data are reported as the number of SHIV RNA copy equivalents per ml of EDTA plasma.

### Measurement of plasma HIV Env-specific IgG by ELISA

HIV Env-specific antibody concentrations in plasma were determined by ELISA (33). Microtiter plates were coated with 1086Δ7 gp120K160N (3 μg/ml) overnight at 4°C and blocked with PBS + 4% whey, 15% normal goat serum, and 0.5% Tween 20. Serially diluted plasma was added to the plate following extensive washing. IgG antibodies were detected with peroxidase-labeled anti-monkey IgG (Southern Biotech), followed by tetramethylbenzidine (TMB; KPL) and stop solution. Absorbance was read at 450 nm immediately after addition of the stop solution. The simianized CD4 binding site monoclonal antibody B12R1 was used as a standard (40). The concentration of HIV Env-specific IgG was calculated using a five-parameter fit curve relative to the standard using SoftMax Pro 6.3 software (Molecular Devices). To account for non-specific binding, the positivity cutoff was selected as the concentration corresponding to 3 times the OD of blank wells.

### Measurement of Env-specific antibodies by binding antibody multiplex assay (BAMA)

Salivary IgG and IgA and plasma IgA antibodies to gp120 were measured using a customized multiplex assay with 1086.cΔ7 gp120-conjugated fluorescent magnetic beads as previously described (33). Prior to performing IgA assays, specimens were depleted of IgG using Protein G Sepharose (GE Healthcare) as described (41). Concentrations of gp120-specific antibodies in saliva were normalized relative to the total IgA or IgG concentrations, which were measured by ELISA. Results for saliva are presented as specific activity (ng anti-gp120 IgA or IgG antibody per μg total IgA or IgG, respectively).

### Antibody avidity

The avidity of IgG antibodies to 1086.cΔ7 gp120, 1086.C V1V2, gp70 consensus C V3 (33), 1157ipd3N4 gp120, and 1157(QNE)Y173H V1V2 was determined using purified total plasma IgG and surface plasmon resonance (SPR) using a Biacore 4000 instrument as described previously (33). The relative avidity score equals the binding response divided by the dissociation rate constant.

### Antibody-dependent cellular cytotoxicity (ADCC)

ADCC activity was measured as previously reported (42). Briefly, CCR5^+^ CEM.NK^R^ T cells (AIDS Reagent Program) were coated with 1086.c or SHIV-1157ipd3N4 gp120 protein. ADCC activity was determined by the GranToxiLux (GTL) assay as described (33, 42, 43). Four-fold serial plasma dilutions beginning at 1:100 were incubated with target cells and human PBMCs from a cryopreserved leukapheresis of an HIV-seronegative donor with the 158F/V genotype for FcγRIIIa after thawing and overnight rest (43-45). ADCC function is reported as endpoint titers determined by interpolation of plasma dilutions that intercepted the positive cutoff and as the maximum proportion of target cells positive for active granzyme B (maximum activity).

### Infected cells antibody binding assay (ICABA)

Plasma antibody binding to HIV-1 Env expressed on surface of infected cells was measured using an infected cell binding assay as previously described (28, 46). Briefly, CEM.NKR_CCR5_ cells were mock-infected or infected with a replication competent infectious molecular clone virus encoding the 1086.c Env (47) for 48-72 hours. Cells were then cultured in the presence of diluted plasma samples from study infants. Cells were subsequently stained with a viability marker, anti-CD4 antibody (clone OKT4, eBiosciences), fixed, and permeabilized prior to staining with a FITC-conjugated goat anti-rhesus IgG (H+L) polyclonal antibody (Southern Biotech). Data represent the frequency of cells positive for IgG-binding to Env for post-vaccination samples compared to the pre-vaccination sample. Values were normalized by subtraction of the frequency of positive cells observed for cells stained with secondary antibody alone and mock-infected cells.

### Antibody-dependent cellular phagocytosis (ADCP)

ADCP assay was performed as previously described (48, 49). HIV envelope (Env) 1086.c K160N gp120 protein was produced in-house by transfection of 293T cells. For ADCP, the HIV Env 1086.c K160N gp120 protein was conjugated to biotin using the Fast Type A Biotin Conjugation kit (Abcam) and then captured on avidin-labeled fluorescent beads (NeutrAvidin^™^, Invitrogen). To form immune complexes with Env-expressing beads, plasma (1:50 dilution), positive antibody controls (HIVIG, RIVIG, VRC01), or irrelevant antibody control (influenza-specific monoclonal antibody, CH65) were incubated with antigen-conjugated beads at 37 ºC for 2 hours. All monoclonal antibody controls were used at a concentration of 25 μg/ml. Immune complexes were then subjected to spinoculation at 1,200 x g in presence of a human-derived monocyte cell line, THP-1 cells (ATCC TIB-201) for 1 hour at 4ºC. Following spinoculation, bead-conjugated antigens and cells were incubated at 37 ºC to allow for phagocytosis to occur. After 1 hour incubation, THP-1 cells were fixed with 2% paraformaldehyde (Sigma) and fluorescence of the cells was assessed by flow cytometry (BD, Fortessa). A “no antibody” control consisting of PBS supplemented with 0.1% bovine serum albumin (1X PBS+0.1% BSA) was used to determine the background phagocytosis activity. Phagocytosis scores were calculated by multiplying the mean fluorescence intensity (MFI) and frequency of bead-positive cells and dividing by the MFI and frequency of bead-positive cells in the PBS/BSA control. All plasma samples were tested in two independent assays and the average phagocytic scores from these 2 independent assays was reported.

### Neutralizing antibody characterization

Neutralizing antibodies were tested as previously reported (50). Briefly, serum was heat-inactivated for 1 hour at 56°C and diluted in cell culture medium and pre-incubated with HIV-1 pseudotyped virus (51) for 1 hour. Following pre-incubation, TZM-bl cells were added and incubated for 48 hours. Cells were subsequently lysed and luciferase activity was determined using a luminometer and BriteLite Plus reagent (PerkinElmer). Neutralization titers were defined as the serum dilution which reduced relative light units by 50% relative to control wells after background subtraction.

### Flow Cytometric Analysis

*T cell activation:* PBMC were isolated from blood as previously described (36). 10^6^ PBMC were stained with surface antibodies listed in Table 2 at room temperature for 20 minutes in the dark. Cells were treated with Cytofix/ Cytoperm (BD Biosciences) per the manufacturer’s protocol and subsequently stained with intracellular marker antibodies (Table 2) in the same manner. Stained cells were fixed with 1% paraformaldehyde (Electron Microscopy Services). 300,000 events were collected using a BD LSRFortessa and analyzed using FlowJo v10.6.1.

**Table 2:**
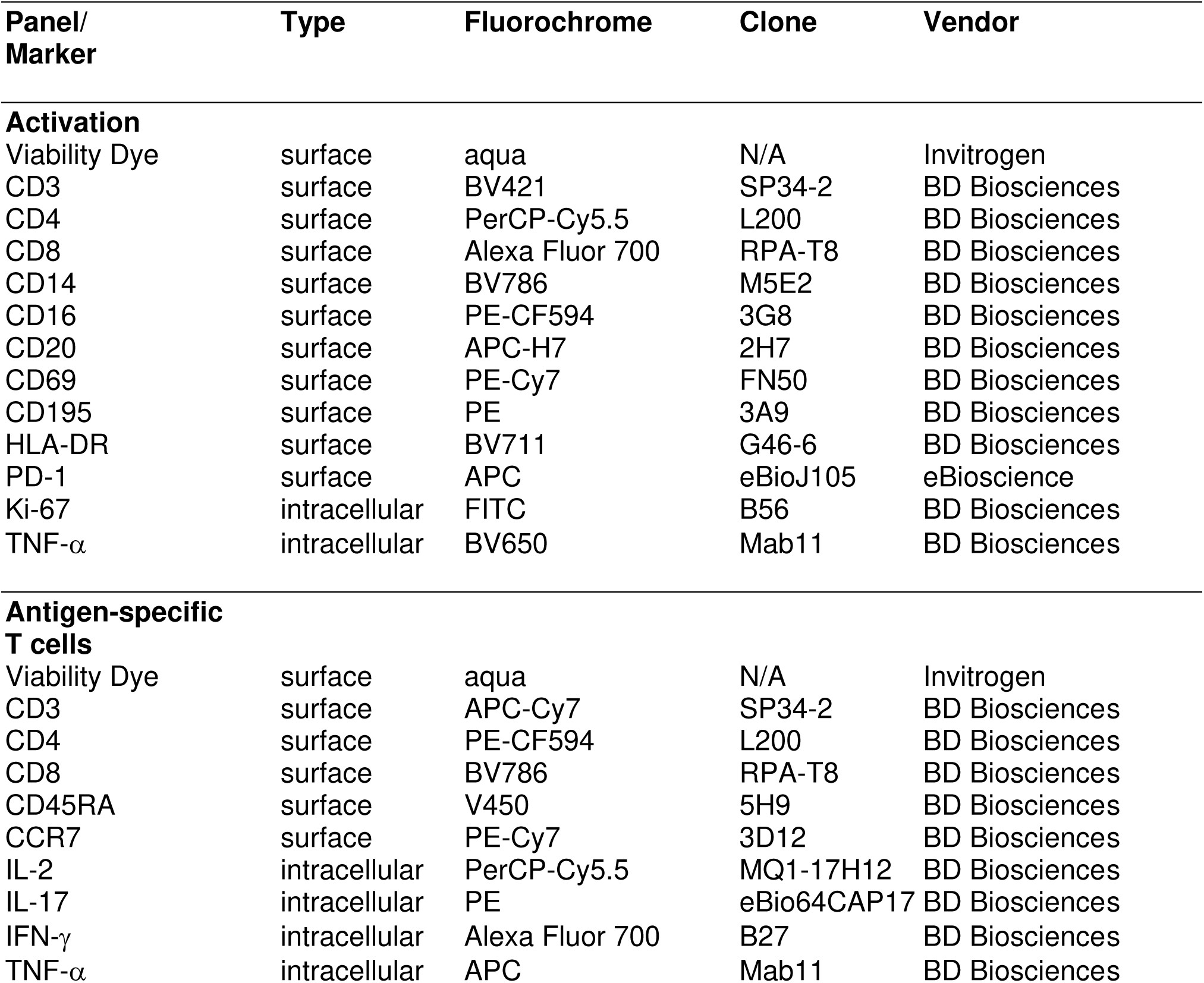
FACS reagent information.

*SIV Gag-specific T cell responses:* SIV Gag-specific T cell responses were determined as described previously (52). Briefly, 2×10^6^ cells were cultured in RPM 1640 supplemented with glutamine, 10% heat-inactivated FBS, and Penicillin/Streptomycin and stimulated with i) vehicle DMSO, ii) 0.5X Cell Stimulation Cocktail (eBiosciences), or iii) 5μg SIV p27Gag peptide pool (NIH AIDS Reagent Program) for 6 h, with 1X Brefeldin A present after the first hour. Cells were stained with antibodies (Table 2) and analyzed as above.

### Statistical Analyses

Statistical tests were performed using R version 3.6.2.

#### Probability of Infection

Kaplan-Meier curves and log-rank tests with exact p-values were used to assess differences between the two groups in the probability of infection at any challenge dose. We presented curves and tested for differences in the probability of infection at any dose. One animal missed seven weekly challenges before resuming challenges on the 1:100 dose and becoming infected on their first 1:100 dose challenge. Thus, the animal was treated as censored at their seventh challenge, (a 1:1000 dose). We estimated the per-challenge probability of infection at each dose administered (1:1000, 1:100, 1:10, 1:2, 1:1) as [# animals infected by a challenge at this dose] / [total number of challenges (across and within all animals) administered up to and including the week of infection at this dose]. For each per-challenge probability of infection, we constructed an approximate 95% confidence interval (Wilson score interval without continuity correction) by assuming that all challenges across and within animals are independent.

#### Antibody correlates of protection

We assessed the association of Env-specific plasma IgG, salivary IgG, salivary IgA, antibody avidity, ADCC, infected cell binding, and ADCP at week 15 with the number of challenges required to achieve SHIV infection in vaccinated animals only. Spearman’s rank correlation coefficients were estimated to assess these associations. All correlations were tested with exact p-values to assess whether any were significantly different from 0. To adjust for multiple comparisons, the Benjamini-Hochberg (BH) procedure was used to control the false discovery rate (FDR). An adjustment to control the FDR at a value of 0.05 was performed for these pre-specified endpoints for a total 16 parameters (Table 3).

**Table 3:** Correlation between Immune Parameters and Number of Challenges to Infection.

| Parameter | N | Spearman Correlation<br>r value | Correlation<br>p value<sup>c</sup> | FDR<sup>a,b</sup><br>p-value | Figure |
| --- | --- | --- | --- | --- | --- |
| **Env-specific IgG<sup>a</sup>** |  |  |  |  |  |
| 1086.c | 12 | -0.519 | 0.0864 | 0.4043 | 2A |
| 1157ipd3N4 | 12 | -0.470 | 0.1246 | 0.4043 | 4A |
| **Plasma IgM<sup>a</sup>** | 11 | -0.288 | 0.3885 | 0.5651 |  |
| **Salivary IgG<sup>a</sup>** | 12 | -0.456 | 0.1373 | 0.4043 | 2C |
| **Salivary IgA<sup>a</sup>** | 12 | -0.189 | 0.5516 | 0.6304 |  |
| **Avidity Index<sup>a</sup>** |  |  |  |  |  |
| 1086.c | 12 | -0.368 | 0.2365 | 0.4553 | 3A |
| 1157ipd3N4 | 12 | -0.165 | 0.6064 | 0.6468 | 4B |
| SHIV1157(QNE)Y175H | 10 | -0.372 | 0.2880 | 0.4608 | 4B |
| 1086.C V1V2 | 11 | -0.119 | 0.7282 | 0.7282 | 3A |
| gp70 ConsC V3 | 12 | -0.428 | 0.1655 | 0.4043 | 3A |
| **ADCC Titer<sup>a</sup>** |  |  |  |  |  |
| 1086.c | 12 | -0.761 | 0.0054 | 0.0870 | 3C |
| 1157ipd3N4 | 12 | -0.568 | 0.0580 | 0.4043 | 4C |
| **ADCC Activity<sup>a</sup>** |  |  |  |  |  |
| 1086.c | 12 | -0.354 | 0.2561 | 0.4553 | 3D |
| 1157ipd3N4 | 12 | -0.418 | 0.1769 | 0.4043 |  |
| **ADCP Score<sup>a</sup>** | 12 | -0.193 | 0.5455 | 0.6304 | 3E |
| **Infected Cell Binding<sup>a</sup>** | 12 | 0.242 | 0.4446 | 0.5928 | 3F |
| **T Cell Activation<sup>b</sup>** |  |  |  |  |  |
| CCR5<sup>+</sup>CD4<sup>+</sup> | 24 | -0.065 | 0.7631 | 0.7648 | 7 |
| Ki67<sup>+</sup>CD4<sup>+</sup> | 24 | -0.161 | 0.4499 | 0.6748 | 7 |
| CD69<sup>+</sup>CD4<sup>+</sup> | 24 | 0.379 | 0.0688 | 0.4131 | 7 |
| PD1<sup>+</sup>CD4<sup>+</sup> | 24 | -0.064 | 0.7648 | 0.7648 | 7 |
| TNF-α<sup>+</sup>CD4<sup>+</sup> | 24 | -0.182 | 0.3913 | 0.6748 | 7 |
| **Env-specific B cells<sup>b</sup>** | 12 | -0.336 | 0.2845 | 0.6748 |  |

<sup>a</sup> or <sup>b</sup> FDR adjustment for multiple comparisons for the sets of tests specified by the subscript
<sup>a</sup> or <sup>b</sup> as described in Materials and Methods
<sup>c</sup> exact p-value to test whether the correlation appears to be significantly different from 0

#### Cellular correlates of protection

Wilcoxon rank-sum tests with exact p-values were used to compare the CCR5^+^ (CD195^+^), Ki-67^+^, CD69^+^, and CD279^+^ (PD1) CD4^+^ T cells at week 15 between vaccinated and control animals. An adjustment to control the FDR at an α value of 0.05 was performed for these 5 endpoints using the BH procedure.

Spearman’s rank correlation coefficients were estimated for the cohort as a whole as well as for vaccinated animals only. All correlations were tested with exact p-values to assess whether any were significantly different from 0. The entire cohort was used to assess whether there was an association between CD4^+^ T cell activation parameters at week 15 and the number of challenges required for SHIV infection. Vaccinated animals were used to assess whether there appeared to be an association between Env-specific B cells and the number of challenges required for SHIV infection. To adjust for multiple comparisons, the Benjamini-Hochberg (BH) procedure was used to control the false discovery rate (FDR). An adjustment to control the FDR at an α value of 0.05 was performed for these pre-specified correlation endpoints for a total 6 parameters (Table 3).

## RESULTS

### Study design

The current study utilized infant RM that were randomly divided into 2 groups of 12 at birth (Table 1; Figure 1). Infant RM in the vaccine group were immunized at week 0 with 2×10^8^ PFU of MVA-HIV 1086.c Env construct and 15 μg 1086.c gp120 protein mixed with 3M-052-SE adjuvant by the IM route. To induce T cell responses, infant vaccinees were also primed with 5×10^10^ vp of ChAdOx1.tSIVconsv239 at week 0. At weeks 6 and 12, infants in the vaccine cohort received IM booster immunizations with MVA-HIV Env, 1086.c gp120 protein in 3M-052-SE, and 2×10^8^ PFU MVA.tSIVconsv239. Control infants received an empty MVA vector at weeks 0, 6, and 12 (Figure 1). Once weekly oral SHIV challenges were initiated at week 15, 3 weeks after the last immunization. Animals were followed for approximately 12 weeks post SHIV-infection, with infection being defined as an animal having two consecutive positive viral RNA results.

### Vaccine-induced 1086.c envelope-specific antibody responses

We first aimed to confirm our prior findings that the vaccine regimen induces potent HIV Env-specific antibody responses (35). Plasma 1086.c gp120-specific IgG responses were detected as early as week 3 after the first immunization in the majority of animals (Figure 2A). Antibody levels were enhanced following the week 6 booster immunization, waned slightly thereafter, and reached peak levels after the final immunization at week 12. Geometric mean plasma HIV Env-specific IgG concentrations at week 14 (1,060401 ng/ml; 95% CI [1,470,184; 21020]) were comparable to those elicited in our prior study (1,251,467 ng/ml; 95% CI [1,049,651; 53481]) (35). We also tested for the induction of Env-specific plasma IgA antibody in vaccinated infants (Figure 2B). The induction of plasma Env-specific plasma IgA was delayed compared to plasma IgG and was of lower magnitude. Env-specific IgG and IgA were also detectable in saliva (Figures 2C, D). The positive correlation between plasma and salivary Env-specific IgG and IgA (Figures 2E, F) implied that antibodies in saliva likely reflected transudation from the plasma rather than local induction at mucosal sites.

**Figure 2:**
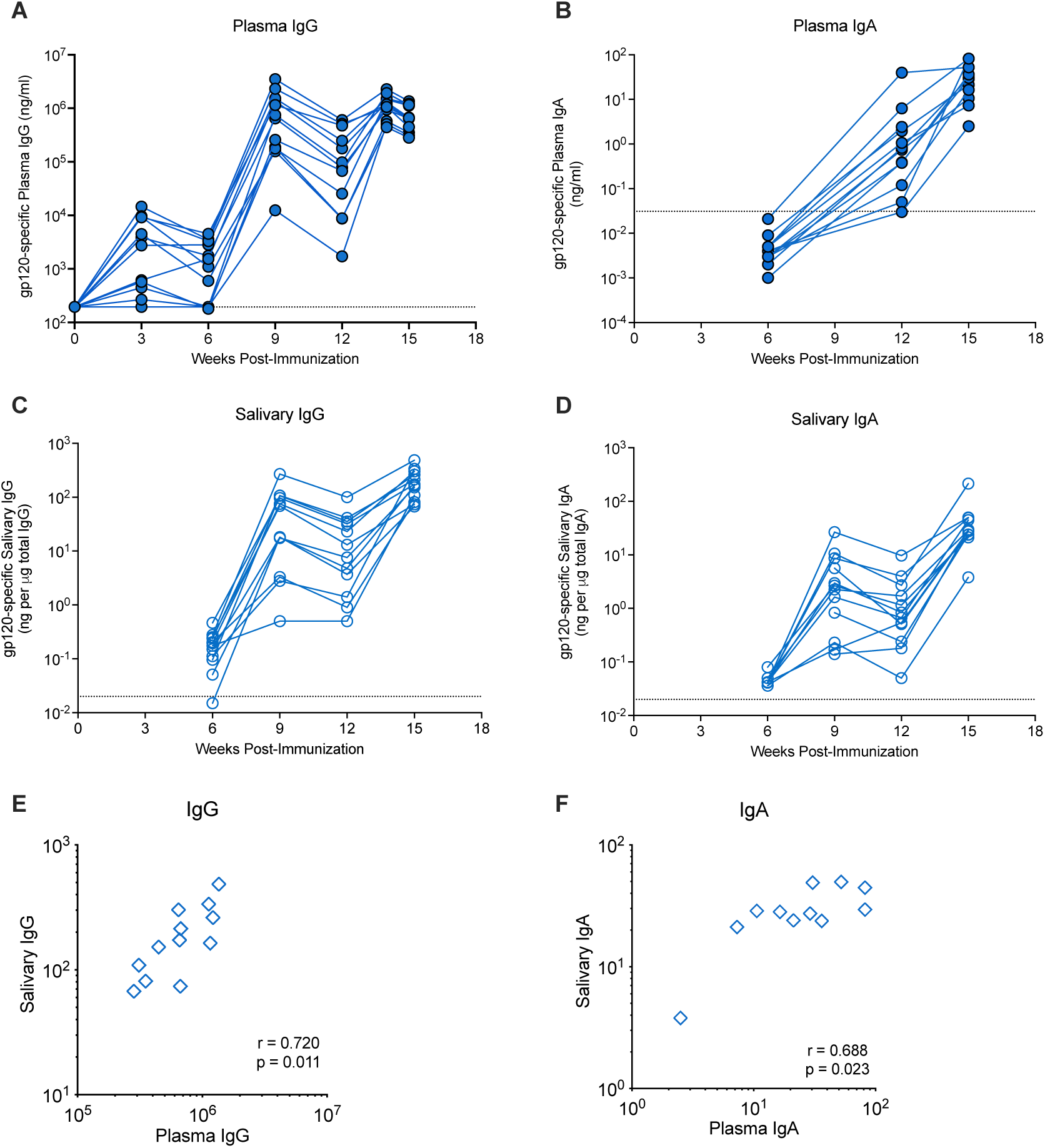
C.1086 Env-specific antibody responses. Plasma (filled blue circles) concentration of 1086.c gp120-specific IgG (Panel A) and IgA (Panel B) were measured by ELISA and BAMA, respectively. Salivary IgG and IgA levels (empty blue circles), measured by BAMA, are reported as specific activity in ng of 1086.c gp120 IgG or IgA per μg of total IgG (Panel C) or IgA (Panel D). Dashed lines represent the cut-off for positivity defined as mean antibody levels in control animals plus 3 standard deviations (SD). Panels E and F illustrate the Spearman correlation between plasma and saliva vaccine-induced IgG or IgA levels, respectively.

We next evaluated the avidity and functional potential of Env-specific plasma IgG. The avidity of plasma IgG specific for 1086.c gp120 was measured by SPR and the median avidity score at week 14 was determined to be 2.4×10^7^ (95% CI [1.45 ×10^7^,6.5 ×10^7^]) (Figure 3A), an avidity similar (p=0.4; Wilcoxon rank-sum test) to the one in our previous study (median avidity score: 4.6×10^7^; 95% CI [1.2×10^7^ to 9.6×10^7^]) (35). The avidity of plasma vaccine-elicited IgG was stronger for the clade C consensus V3 compared to the V1V2 epitope of 1086.c Env (Figure 3A). The current vaccine regimen elicited weak clade C Tier 1 neutralization antibodies. In 7 of 12 vaccinated infants, the peak neutralization titers against the Tier 1b virus I6644.v2.c33 were >500 at week 14, but only 5 of the 7 animals had maintained Tier 1b ID_50_ titers >500 by week 15 (Figure 3B).

**Figure 3:**
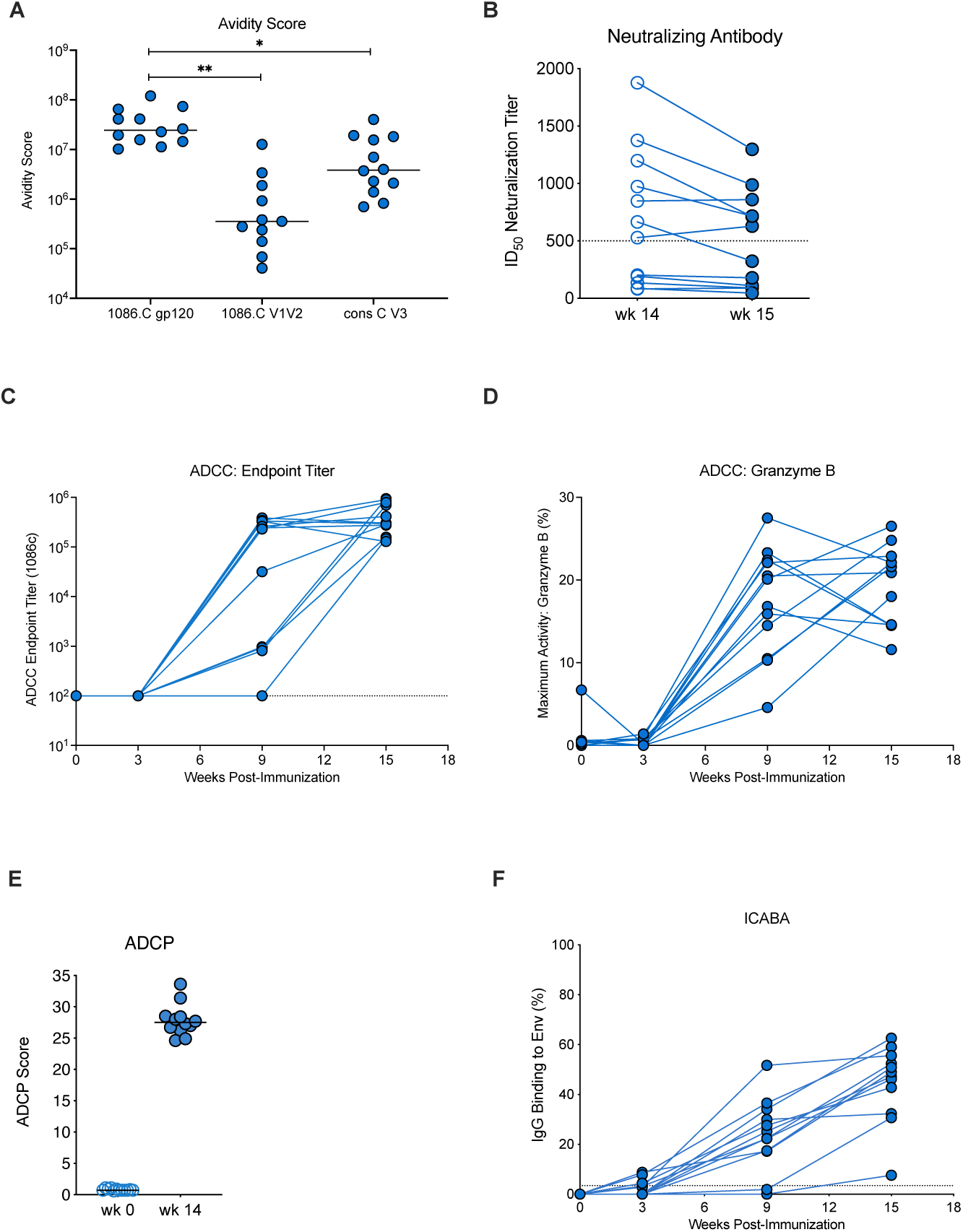
Pre-challenge antibody function of vaccinated infant macaques. Panel A: Avidity Score, determined by SPR, of week 15 plasma IgG specific for 1086.c gp120 or V1V2, or for the consensus clade C V3 (gp70). Each symbol represents a single animal. Panel B: Tier 1b clade C I6644.v2.c33 neutralization titers of vaccinated infants at weeks 14 (empty circles) and week 15 (filled circles). Panels C and D: Longitudinal data for ADCC endpoint titers and maximum granzyme B activity, with each line representing an individual animal. Dashed lines indicate the limit of detection. Panel E: ADCP Scores for vaccinated animals prior to vaccination at week 0 (empty circles) and week 14 (filled circles). Panel F: The ability of plasma IgG binding to cells infected with HIV 1086.c are shown over time for individual vaccinated animals.

Because a main goal of the current design was to elicit non-neutralizing antibody, we assessed the propensity of vaccine-elicited plasma antibody for FcR-mediated ADCC. ADCC responses against Env 1086.c gp120 were detectable in 75% of vaccinated infants by week 9 and in 100% by the time of initial SHIV challenge at week 15 (Figure 3C). Similar to vaccine-induced plasma Env-specific IgG, the high ADCC endpoint titers and median granzyme B activity (Figure 3D) in the current study were comparable to those observed in our prior study (35). In addition to ADCC, 1086.c Env-specific plasma antibodies were also able to mediate ADCP (Figure 3E). Relevant to both ADCC and ADCP function, plasma IgG was capable of binding to Env 1086.c expressed on the surface of HIV-infected cells (ICABA), with 11 of 12 infants having >20% binding at week 15 (range: 7.60%-62.62%; mean: 44.70%) (Figure 3F).

We also measured antibody responses relevant to the heterologous SHIV challenge virus, including clade C 1157ipd3N4 Env-specific IgG and 1157(QNE)Y173H Env V1V2-specific antibody responses. Although the overall magnitude of plasma binding antibodies to 1157ipd3N4 gp120 was lower compared to 1086.c gp120-specific IgG, the kinetics of plasma binding antibodies to 1157ipd3N4 Env followed a similar pattern as observed for 1086.c gp120-specific IgG. All animals developed 1157ipd3N4 gp120-specific IgG after the second immunization with peak responses at week 14, two weeks after the third immunization (Figure 4A). The median avidity score of plasma IgG against 1157id3N4 gp120 (1.9×10^6^; 95% CI [7.3 ×10^6^, 2.3 ×10^6^]) was about 1 log lower compared to the avidity index for the vaccine immunogen 1086.c gp120, and the avidity for the V1V2 region of 1157(QNE)Y173H was one log lower compared to the avidity for 1086.c V1V2 (Figures 4B and 3A). The vaccine regimen elicited high levels of Env-specific plasma antibodies with ADCC activity against the clade C 1157ipd3N4 gp120 (Figure 4C), with median endpoint titers (2.89×10^5^) comparable to the median titer for 1086.c (2.99×10^5^; Figure 3B), although ADCC 1157ipd3N4-specific IgG endpoint titers exhibited greater variability among the individual animals.

**Figure 4:**
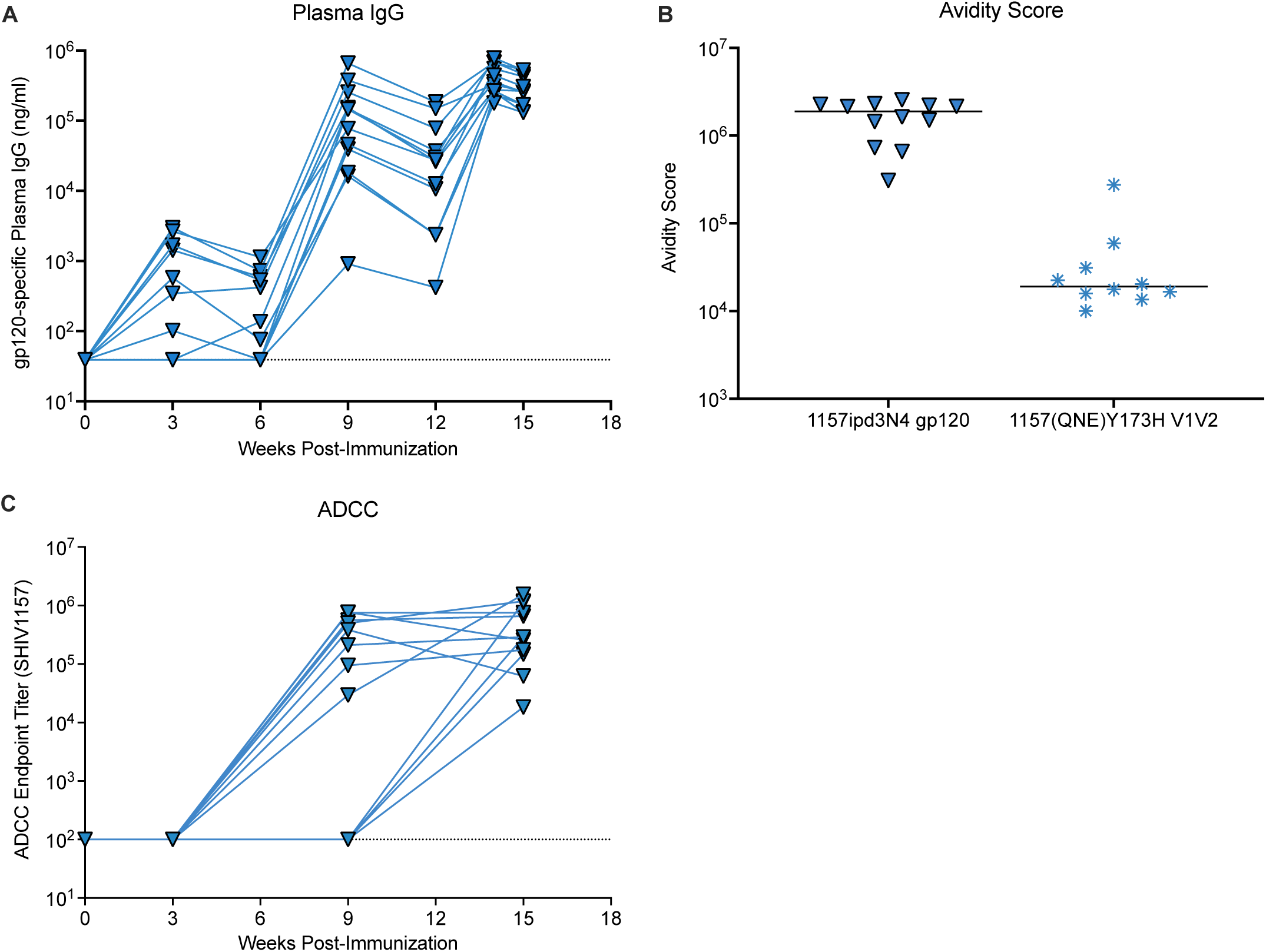
Vaccine-induced 1157ipd3N4 and SHIV1157(QNE)Y175H Env-specific antibody responses. Panel A: Plasma concentration of 1157ipd3N4 gp120-specific IgG (blue triangles) over time. Panel B: Avidity scores of plasma IgG specific for 1157ipd3N4 gp120 (blue triangles) or gp70-V1V2 SHIV1157(QNE)Y375H (blue star symbols). Each symbol represents an individual animal; horizontal lines represent the median. Panel C: ADCC endpoint titers for plasma antibodies specific to 1157ipd3N4 gp120 (blue triangles).

### Cellular responses to vaccination

The majority of vaccinated animals developed SIV Gag-specific T cell responses by week 14 in peripheral blood (Figure 5A). In lymph nodes, however, SIV Gag-specific T cell responses were only detected in 9 of 12 vaccinees (Figure 5B). SIV Gag-specific CD4^+^ T cells appeared to produce predominantly TNF-α and IL-17, whereas a more mixed cytokine response was observed in CD8^+^ T cells. Polyfunctional cytokine responses were rare.

**Figure 5:**
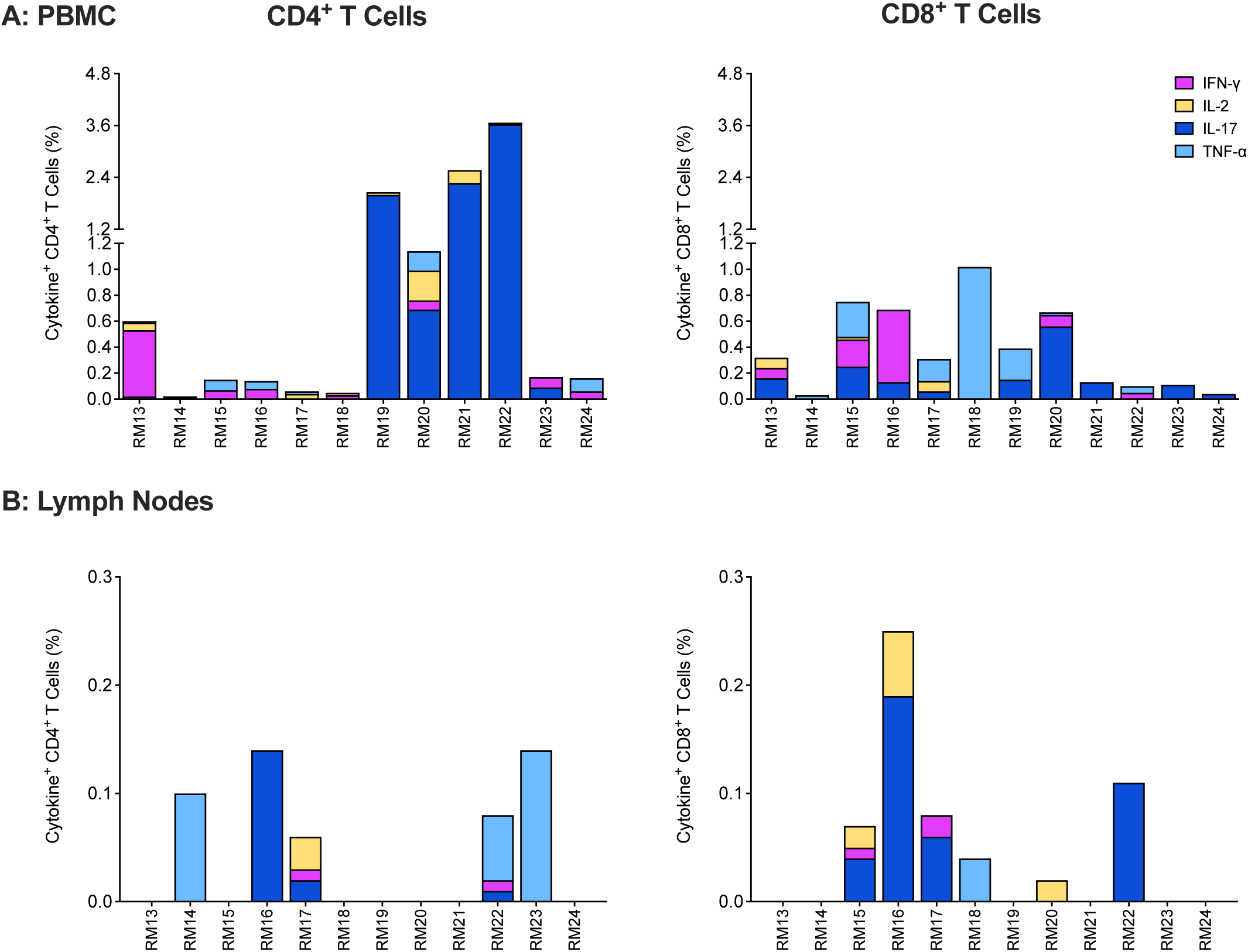
SIV Gag-specific T cell responses in PBMC and peripheral lymph nodes at week 14. Each bar in Panels A and B represents the sum of single cytokine responses of SIV Gag-specific CD4^+^ (left graphs) or CD8^+^ T cells (right graphs) for each vaccinated animal at week 14 in PBMC (Panel A) or lymph nodes (Panel B). Cytokines measured include IFN-γ (fuchsia), IL-2 (yellow), IL-17 (dark blue), and TNF-α (light blue).

### SHIV1157(QNE)Y173H challenge outcome

Starting at week 15, 3 weeks after the third immunization, animals were challenged once weekly with SHIV by the oral route. The initial virus dose consisted of 1:1,000 diluted virus stock, a dose that was purposely chosen to be 10-fold higher compared to the dose (1:10,000) successfully used in an intrarectal challenge study in adult rhesus macaques (28), because of the lower risk estimate for oral versus IR infection determined by human HIV epidemiologic studies (53) and SHIV1157 infections in adult RM (54). Seven of 12 control vaccinated infants became infected at the 1:1000 SHIV dose and 4 of the remaining 5 animals became infected at 1:100. RM10 remained uninfected after 29 challenges and only became infected after oral challenge with undiluted viral stock (Figure 6A, Table 1). The median challenge number required to become infected for control vaccinated infants was 7.5. In comparison, vaccinated animals required a median number of 11 challenges to achieve infection (Figure 6B). Half of the vaccinated animals (n=6) were infected at the 1:1000 dose and three additional animals by the 1:100 dose. The remaining 3 animals were infected by 1:10 (n=2) and 1:2 (n=1) challenge virus dilutions (Figure 6A). Although vaccinated animals required a slightly higher average number of challenges to infection (11 exposures) compared to controls (7.5 exposures), there was no difference in the probability of infection at any challenge dose between the two groups (p=0.89; Figure 6C). When we compared the probability of infection between control and vaccinated animals that became infected at the 1:1,000 challenge virus dose, at the 1:1000 or 1:100 dose, or at the 1:1000, 1:100, or 1:10 doses, we also did not detect differences in infection risks. The distribution of peak viremia also did not differ between vaccinated animals (median: 1.85×10^7^ viral RNA copies/ ml) and control animals (median: 7.5×10^6^ viral RNA copies/ ml; Wilcoxon rank-sum test with exact p=0.24) (Figure 6D).

**Figure 6:**
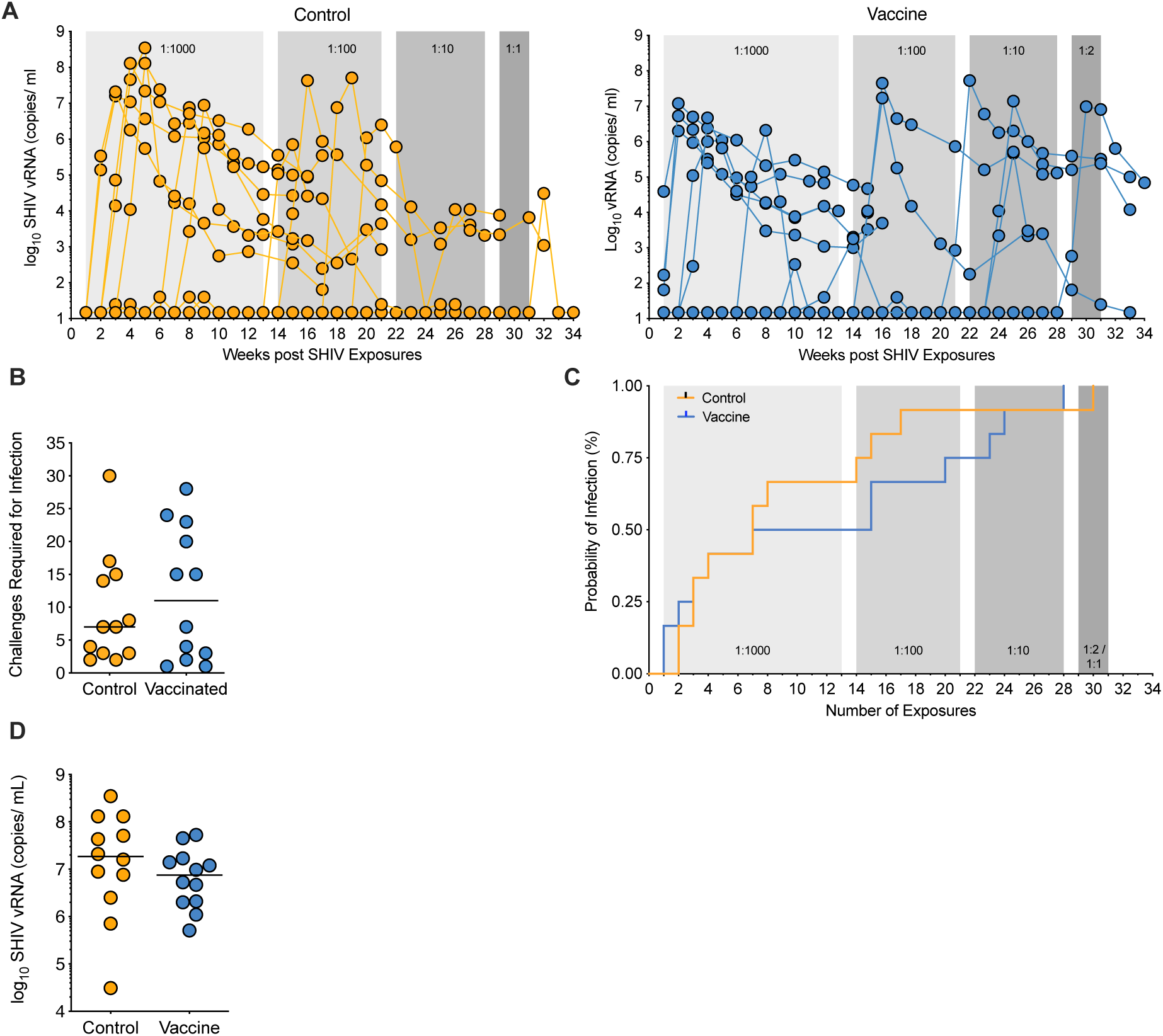
Challenge outcome. Panel A: Longitudinal plasma viral load as assessed by RT-PCR from control and vaccinated cohorts of infant RM are displayed. Shaded areas represent the challenge dose: light gray, 1:000, weeks 0-13; medium gray, 1:100, weeks 14-21; dark gray, 1:10, weeks 22-28; darkest gray, 1:2 or undiluted. Panel B: The number of challenges required for infection is plotted for control and vaccinated animals. Horizontal lines represent the median. Panel C: Kaplan-Meier survival curves for any dose of viral stock dilutions are shown for control and vaccinated infants. Panel D: Peak viremia in control and vaccinated animals. Control and vaccinated animals are indicated by orange or blue lines/symbols, respectively, with each symbol representing an individual animal; horizontal lines indicate the median.

These data implied that the vaccine-induced Fc-mediated effector functions of Env-specific plasma IgG were insufficient to protect against oral SHIV challenge in infant macaques. In fact, vaccine virus-specific ADCC activity was associated with fewer challenges required for infection (r= -0.761, unadjusted p=0.0054; FDR adjusted p=0.0870; Table 3; Figure 7A). This trend was most apparent when vaccinated infant RM were stratified by median ADCC titer (2.99×10^5^). The results suggested that animals with ADCC titers below the median required more SHIV exposures to become infected compared to animals with ADCC titers above the median (Figure 7B). However, the probability to infection was not different between control animals, vaccinated animals with ADCC titers below the median, and vaccinated animals with ADCC titers above the median (log rank test with exact p=0.06). Consistent with comparable median ADCC titers for 1086.c and 1157ipd3N4 Env, 1157ipd3N4 Env-specific ADCC titers trended towards a negative correlation with the number of challenges required for infection, although this trend was not substantiated after adjusting for FDR (Table 3; r= -0.568, unadjusted p=0.0580; FDR adjusted for p=0.4043). There was no association between ADCC activity, as assessed by maximum granzyme B production, and challenge outcome. Similarly, neither plasma or salivary Env-specific binding antibodies at week 15, nor the avidity of Env-specific antibodies correlated with challenge outcome (Table 3).

**Figure 7:**
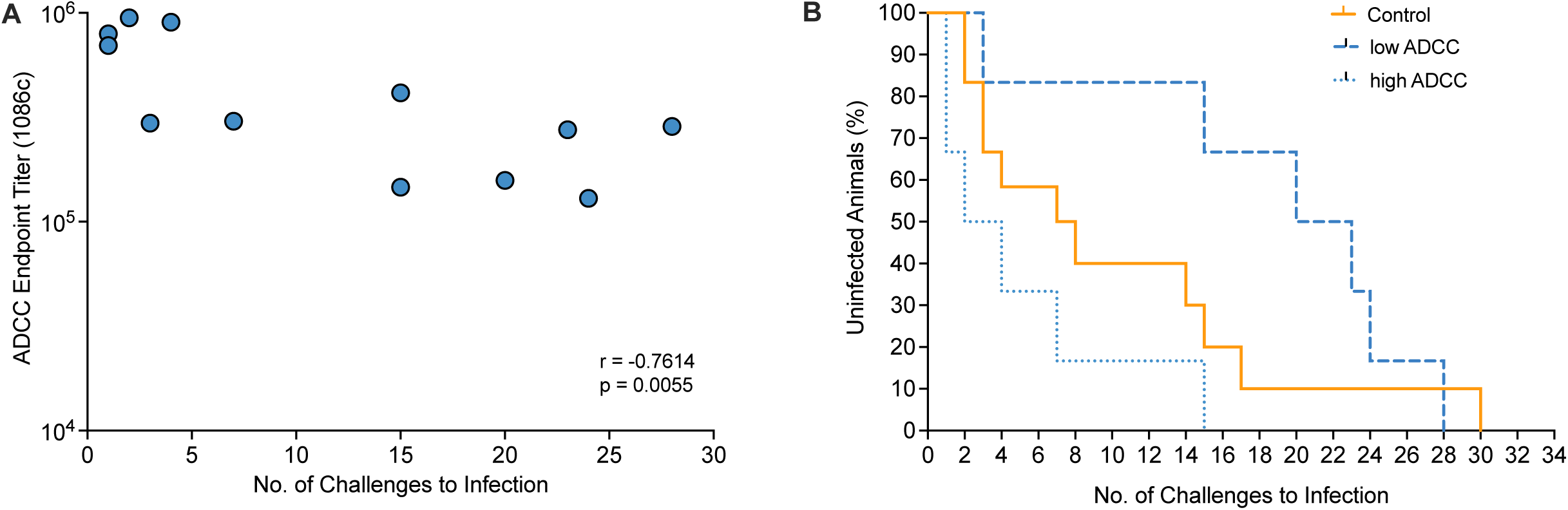
Correlation between ADCC endpoint titers and challenge outcome. Panel A: Graph of the Spearman rank correlation between ADCC endpoint titers and number of challenges required for infection of each vaccinated animal. Panel B: Kaplan Meier plot to demonstrate the relationship between ADCC endpoint titers and number of challenges required for infection when vaccinated animals are categorized as having a low (blue dashed line) or high (dotted blue line) ADCC titer based on the median ADCC endpoint titer of 10^5^ in comparison to control animals (orange line). Mantel-Cox log rank test was applied to determine differences in the risk of infection between groups.

To explore whether the vaccine had caused non-specific immune activation that could promote increased susceptibility to infection (55, 56), we tested for activation of peripheral blood CD4^+^ T cells, the main target cells for HIV. At the time of challenge initiation (week 15) we noted no difference in the frequency distributions of CCR5^+^ (CD195^+^), Ki-67^+^, CD69^+^, or CD279^+^ (PD1) CD4^+^ T cells in blood of vaccinated compared to control animals (Figure 8). Although vaccinated animals had greater median frequencies of PD-1-positive and TNF-α-producing CD4^+^ T cells compared to the control group (Figure 8), there was no correlation with this response and the number of exposures required to achieve infection (Table 3).

**Figure 8:**
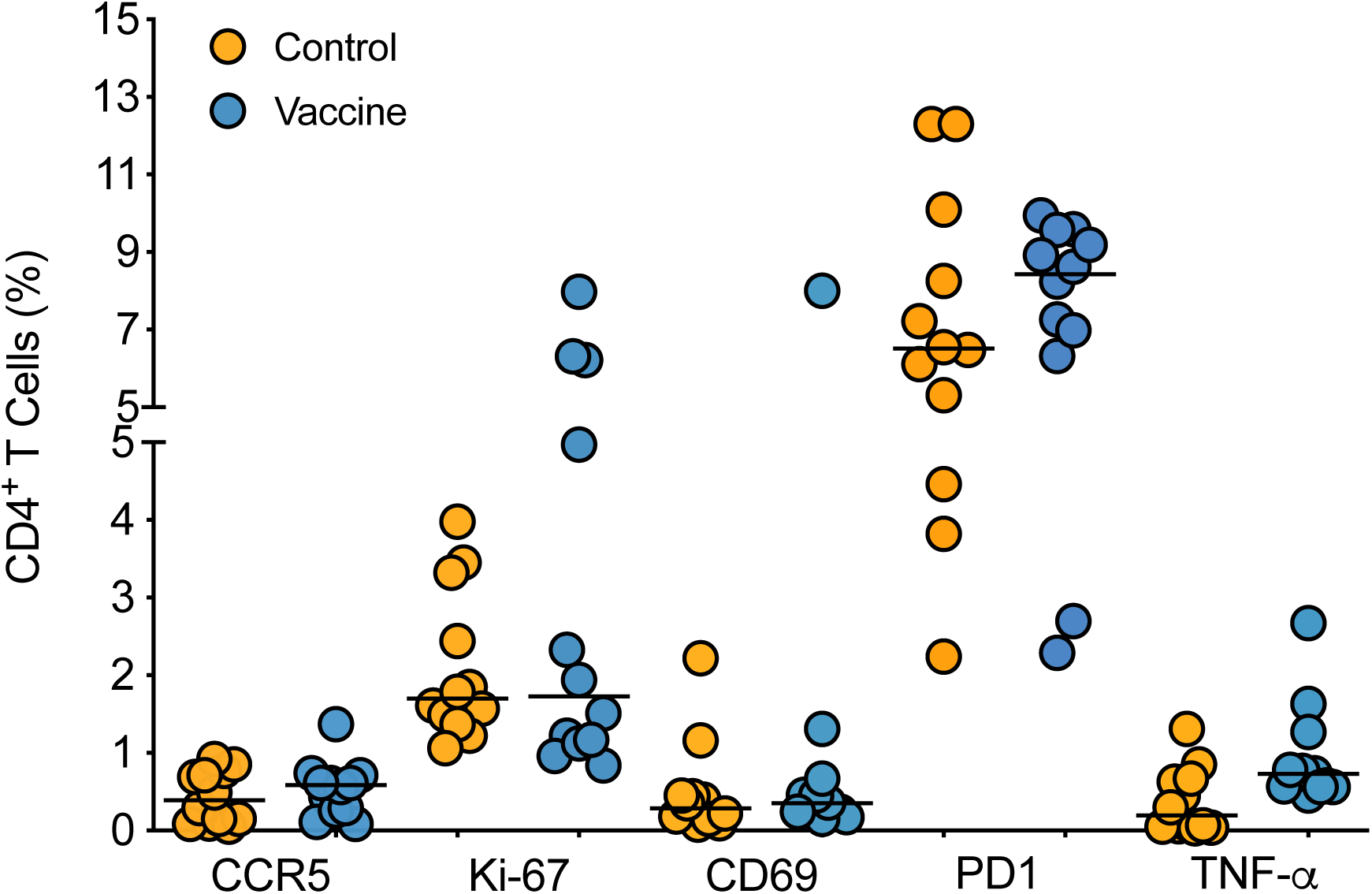
CD4^+^ T cell activation. PBMC from week 15 after vaccination were gated on CD3^+^CD4^+^ T cells and assessed for surface expression of CD195 (CCR5), CD69, and CD279 (PD1), and intracellular expression of Ki-67 and TNF-α. TNF-α positive T cell frequencies between control (orange circles) and vaccinated (blue circles) animals were compared by Mann-Whitney test.

## DISCUSSION

According to the UNAIDS 2021 estimates, in 2020, every day more than 400 children became infected with HIV (1). Therefore, despite increasing access to ART, vaccine development remains an urgent task to prevent new pediatric HIV infections. The current study tested the efficacy of an MVA-Env plus Env protein with 3M-052-SE adjuvant vaccine regimen combined with an ChAdOx.1.tconsSIVma239-Gag, Pol prime, MVA.tconsSIVma239-Gag, Pol boost regimen that had been optimized to maximize Env-specific antibody responses with Fc-mediated effector function (33-35, 42) in infant RM. Several previous HIV vaccine studies in NHP had found a correlation between reduced infection or control of viral replication and vaccine-induced antibodies mediating ADCC (28, 57-59) and/or ADCP and antibody-dependent neutrophil phagocytosis (25-27). However, despite the induction of robust Env-specific antibodies with Fc-mediated effector functions, infant RM receiving the above described vaccine regimen were not protected against oral SHIV infection. There was also no evidence of virus control, a clinically important secondary read-out of vaccine efficacy pertaining to less severe disease outcomes and reduced HIV transmission risk (60).

The reasons for lack of efficacy are likely multifold. We used a challenge virus with an Env that was heterologous to the vaccine immunogen and started challenges shortly (3 weeks) after the last vaccine immunization to closely mimic consistent, real-world exposure of infants breastfed by HIV-infected women. It is possible that some residual activation in response to immunization was still lingering. We (55, 56) and others (61-64) had previously reported that T cell activation can contribute to an enhanced risk of infection with HIV, SIV or SHIV. Although we observed higher frequencies of TNF-α-positive peripheral blood CD4^+^ T cells at the time of the first challenge in vaccinated compared to control animals, T cell activation was not correlated with the number of SHIV challenges required for infection.

In our studies leading up to the current vaccine study (33-35, 42, 56), we had focused on the optimization of Fc-mediated Env-specific IgG responses. Our vaccine regimen was not designed to induce Tier 2 neutralizing antibodies that are thought to be essential in the protection against SHIV infection in RM (65). We had further reasoned that the inclusion of ChAd- and MVA-vectored vaccines expressing SIV Gag, Pol would induce antiviral T cell responses capable of controlling virus replication at the entry site. However, SIV Gag-specific T cell responses elicited by the ChAd- and MVA-vectored vaccines were of relatively low magnitude and neither PBMC nor lymph node CD4^+^ and CD8^+^ T cell responses at week 14 correlated with the number of challenges to infection or with peak viremia.

Our challenge outcome results are consistent with other infant and adult NHP studies that failed to demonstrate efficacy against SIV or SHIV infection by antibodies with Fc-mediated effector function only (65-67) and human HIV vaccine trials following and building on the results of the RV144 trial did not observe a reduced HIV infection risk. In the RV144 trial protective ADCC function was primarily associated with V1V2- and C1-specific antibodies (68, 69). Our vaccine regimen, however, appears to be biased towards the induction of V3 over V1V2-specific and C5 versus C1-specific epitopes (35). Furthermore, plasma IgG responses specific to the V1V2 region of the vaccine 1086.c Env and of the challenge virus SHIV1157(QNE)Y173H were of lower avidity compared to the relevant gp120-specific IgG. Limited plasma volumes prevented us from assessing ADCC and ADCP activity of epitope-specific antibodies in addition to gp120-specific antibodies in the current study. In future studies, more targeted, epitope-specific analyses - including impact of glycosylation and epitope conformation - may prove beneficial in the interpretation of vaccine outcomes (69, 70).

It is also important to note that the detailed analysis of RV144 results found that trial participants with medium levels of ADCC activity had reduced infection risk when compared to participants with low levels of ADCC activity, while there was no such difference when comparing those with high and low vaccine-induced ADCC responses (see supplement of (68)). In the current study, 1086.c-specific plasma antibodies with ADCC activity could be detected at a median endpoint titer of 1:10^5^ at the time of challenge initiation. Paradoxically, although individual animals with ADCC titers >1:10^5^ were as likely to acquire infection as their control counterparts, high 1086.c ADCC titers >1:10^5^ appeared to be associated with fewer challenges to infection. One potential explanation for this observation is an *in vivo* prozone, a phenomenon when high antibody in the presence of limiting antigen results in smaller immune complexes that cluster fewer Fc domain receptors on the surface of target cells and limit killing activity (71, 72). A prozone effect was also described in an early HIV infection study (73) in which plasma IgG concentrations above 10 μg/ml inhibited NK cell lysis. This data, and data from passive immunization of mice (74), suggest that there may be an optimal level, with lower and upper limits, at which non-neutralizing antibodies are most effective. However, what these levels are in the context of different exposures and how they potentially impact challenge outcome is not yet known.

Similarly, it is difficult to discern from the current literature whether there is an optimal ADCP score. Despite several studies suggesting a correlation between ADCP function and reduced HIV risk in human adults (75, 76) or SHIV infection in adult RM (25), ADCP activity elicited by the vaccine tested in the current study was not correlated with protection against oral SHIV1157(QNE)Y375H infection in infant RM. While the simple comparison of various antibody functions across different vaccine regimens, age groups, and challenge regimens is likely flawed, and different assay conditions may further impact data, the results of our study imply that the magnitude of ADCC or ADCP activity alone is not a reliable predictor of vaccine efficacy. More research is needed to assess the impact of antibody subtype, effector cell and specific Fc receptors mediating the specific functions on vaccine efficacy in preclinical NHP studies (77) and how these findings translate to humans (78). Such findings would likely result in improved *in vitro* assays to measure antibody function and thereby enhance the predictive value of these assays for vaccine efficacy assessment. Highly relevant for pediatric studies, age-dependent differences in immune function of effector cells are not considered. There are numerous studies documenting that NK cells and monocytes exhibit reduced functional capacity, including ADCC (79) and phagocytosis, in infants compared to adults (see reviews by: (80-84). Few studies have examined the expression of FcRγI, FcRγII, and FcRγIII on infant NK cells, monocytes, and neutrophils (85, 86). Therefore, in future studies, we will expand the analysis of vaccine-induced B and T cell responses and also determine whether and how pediatric HIV vaccine regimens impact innate immune cells and their functions.

In summary, while the pre-challenge immunogenicity data demonstrated high magnitude effector antibody functions previously tied to some HIV vaccine efficacy, our results imply that Env-specific ADCC and ADCP responses induced by this candidate vaccine regimen were not sufficient to prevent infection with oral tier 2 SHIV1157(QNE)Y375H in infant RM. Therefore, future studies of interventions to protect infants against HIV acquisition through breastfeeding should focus at improving the breadth of the antibody response, namely the induction of bnAbs or passive administration of combinations of long-acting HIV bnAbs, as well as overcoming the relative paucity of cell-mediated immunity induced by current vaccine platforms in early life.

## ACKNOWLEDGEMENTS

The work was supported by National Institutes of Health grants 1R56 DE026321 (KDP), P01 AI117915 (SP, KDP), T32 5108303 (ADC), T32AI007392-31 (SJB), the Office of Research Infrastructure Programs/OD P51OD011107 (to CNPRC), and the Center for AIDS Research award P30AI050410 (to UNC). The UNC Flow Cytometry Core Facility is supported in part by P30 CA016086 Cancer Center Core Support Grant to the UNC Lineberger Comprehensive Cancer Center. Research reported in this publication was supported by the Center for AIDS Research award number 5P30AI050410. The content is solely the responsibility of the authors and does not necessarily represent the official views of the National Institutes of Health.

We thank Sampa Santra (Harvard University) for kindly providing the SHIV-1157(QNE)Y173H viral stock, Dr. Tomáš Hanke (Oxford University, Oxford, UK) for the ChAdOx1.tSIVconsv239 and MVA.tSIVconsv239 vaccines, and IDRI for 3M-052-SE

We are grateful for technical assistance by the CNPRC staff, Jennifer Watanabe, Jodie Usachenko, Amir Ardeshir (CNPRC at UCD), Robert L. Wilson (LSUHSC), R. Whitney Edwards, Nicole Rodgers (Duke University), Neelima Choudary, and Ryan H. Tuck (UNC). The authors also acknowledge Celia LaBranche and colleagues (Duke University) for performing neutralization assays, and David J. Pickup (Duke University) for development and production of the recombinant MVA-based vaccine.

We thank Dr. J. Lifson, Rebecca Shoemaker and their colleagues in the Quantitative Molecular Diagnostics Core of the AIDS and Cancer Virus Program of the Frederick National Laboratory for expert assistance with viral load measurements.

